# Bacterial pore-forming proteins induce non-monotonic dynamics due to lipid ejection and crowding

**DOI:** 10.1101/2020.11.11.378638

**Authors:** Ilanila Ilangumaran Ponmalar, K. G. Ayappa, J. K. Basu

## Abstract

Developing alternate strategies against pore forming toxin (PFT) mediated bacterial virulence factors require an understanding of the target cellular response to combat rising antimicrobial resistance. Membrane-bound protein complexes involving PFTs, released by virulent bacteria are known to form pores leading to host cell lysis. However, membrane disruption and related lipid mediated active repair processes during attack by PFTs remain largely unexplored. We report counter intuitive and non-monotonic variations in lipid diffusion, measured using confocal fluorescence correlation spectroscopy, due to interplay of lipid ejection and crowding by membrane bound oligomers of a prototypical cholesterol dependent cytolysin, Listeriolysin O (LLO). The observed protein concentration dependent dynamical cross-over is correlated with transitions of LLO oligomeric state populations from rings to arc-like pore complexes, predicted using a proposed two-state free area based diffusion model. At low PFT concentrations, a hitherto unexplored regime of increased lipid diffusivity is attributed to lipid ejection events due to a preponderance of ring-like pore states. At higher protein concentrations where membrane inserted arc-like pores dominate, lipid ejection is less efficient and the ensuing crowding results in a lowering of lipid diffusion. These variations in lipid dynamics are corroborated by macroscopic rheological response measurements of PFT bound vesicles. Our study correlates PFT oligomeric state transitions, membrane remodelling and mechanical property variations, providing unique insights into developing strategies to combat virulent bacterial pathogens responsible for several infectious diseases.

**SIGNIFICANCE:** Developing alternate strategies against pore forming toxin (PFT) mediated bacterial virulence factors requires understanding target cellular responses and cellular defence strategies to combat rising antimicrobial resistant strains. While it is well understood that PFTs exist in a wide variety of oligomeric states, the underlying membrane response to these states is unexplored. Using confocal fluorescence correlation spectroscopy and a membrane free area based model we relate non-monotonic variations in the lipid diffusivity arising from an interplay of lipid ejection events and membrane crowding due to variations in concentration of membrane bound listeriolysin O. Our observations have a direct bearing on understanding cellular defense and repair mechanisms effective during initial stages of bacterial infection and intrinsically connected to the underlying membrane fluidity.

## INTRODUCTION

A large class of bacterial pathogens infect target cells by releasing pore forming membrane disruptive proteins whose sole purpose is to compromise metabolic and signalling pathways that often lead to cell lysis (1). Pore-forming toxins (PFTs), a key class of membrane excision causing proteins, are secreted by pathogenic bacteria as a virulence factor for attacking the host or as a defense mechanism against the host immune system (2). The prevalent mechanism of PFT action occurs by forming transmembrane pores on the protective cell membrane leading to cell lysis (1). Listeriolysin O (LLO) which is the focus of this study, comes under the subclass of cholesterol dependent cytolysins (CDCs) secreted by gram positive bacterium *Listeria monocytogenes* implicated in listeriosis, fatal to immune-compromised individuals (3). Recent studies using high resolution cryo-electron microscopy (cryo-EM) (4) and high speed atomic force microscopy (AFM) (5) of pore formation by CDCs on model membranes have revealed the kinetics of pore formation by including existence of pre-pore and pore states and the presence of a distribution of oligomeric states resembling arcs and rings (5–7). While it is known that the nature of these oligomeric states of PFTs depend on membrane fluidity (5, 8, 9) and, in case of CDCs, on availability of cholesterol (10), the manner in which incorporation of these states influences biomembrane dynamics is unclear. Membrane fluidity is also believed to play an important role in cellular repair processes such as exocytosis and endocytosis (11).

In case of membrane excision causing proteins that alter lipid diffusion such as antibiotics (12, 13), pore-forming toxins (8, 9, 14–16), and peptides (17–19), it is not clear whether existing models for integral membrane protein concentration dependent membrane dynamics will be applicable (20). A problem central to PFT attack is the manner in which lipids are either displaced or ejected during pore formation. Recent molecular dynamics simulations reveal that lipid ejection (21, 22) can occur during insertion of fully oligomerized pores or rings and lipids are displaced to stabilize membrane inserted arcs (21, 23). To our knowledge, the only direct evidence for lipid ejection, is from AFM experiments with the CDC, suilysin (24). Processes such as lipid ejection and displacement are expected to influence the available free area for lipid mobility in a concentration dependent manner. Proteins can also influence membrane mechanical properties such as bending and stiffness moduli which influence the propensity of cells to lyse (25–27). However little is known about how these macroscopic changes in the mechanical and rheological properties influence membrane repair processes when exposed to PFTs (28). Our earlier studies have revealed a strong connection between lipid dynamics measured using confocal and super resolution stimulated emission depletion (STED) fluorescence correlation spectroscopy (FCS) and the efficacy of pore formation by LLO and cytolysin A (Cly A), an *α*-PFT, dependent on the specific phase of the lipid membranes (8, 9, 15). Recent experiments with GUVs exposed to LLO, enabled correlating changes in lipid dynamics to membrane bound conformational states of LLO oligomers (8), which was also found to depend on LLO concentration. Additionally repair mechanisms in cells to alleviate CDC attack are active when pores form at lower protein-to-lipid ratios where effective toxin removal mechanisms have been observed (29). Hence a concentration dependent study of interaction of PFTs with biomembranes can help unravel the interplay between membrane excision and repair mechanisms intrinsically connected to membrane fluidity and protein mediated host cellular trafficking and signalling events.

In this article, we report experiments on supported lipid bilayers exposed to LLO and monitor lipid dynamics and membrane micro-rheology over a broad range of protein-to-lipid ratios encompassing the dilute, intermediate as well as the crowded regime. This systematic variation of protein concentration reveals a unique non-monotonic behaviour of lipid diffusivity as a function of PFT concentration which falls outside the purview of existing membrane dynamics models. Not only do we observe the anticipated crowding induced decreasing lipid diffusivities at high protein concentration, but our results reveal a counter intuitive increase in lipid diffusivity at intermediate LLO concentrations. We quantitatively capture these trends using an established model based on the dependence of membrane lipid diffusion (30) on the available free area. We correlate changes in observed lipid diffusivity to the free area contributions from different PFT oligomeric states, thus providing unique insights connecting lipid dynamics to bound protein distributions.

We ascribe the increasing diffusivity regime to active membrane remodeling and lipid ejection events, predominantly due to ring-like pore formation, increasing available free area and enhancing lipid mobility. Interestingly, the decreasing diffusivity regime arises from a preponderance of arc-like structures which co-exist with the ring-like pores, but are not as efficient in lipid ejection, as reported earlier (21, 22). This combined effects of lower lipid ejection efficiency by arcs and reduction in available free area akin to crowding effects due to increasing protein concentration leads to the observed decrease in lipid diffusivity (17, 31–33). These regimes are also qualitatively captured in the non-monotonic trends observed in the loss modulus of small unilamellar vesicles exposed to proteins in diffusing wave spectroscopy (DWS) micro-rheology experiments. These results bring forth a unique correlation between PFT oligomeric state transitions and membrane dynamics coupled with mechanical properties which can provide insight into developing effective strategies to combat these virulent pathogens responsible for causing several infectious diseases.

## MATERIALS AND METHODS

### Listeriolysin O purification, labelling and lytic activity

Point mutations were made on pPROb LLO plasmid using site-directed mutagenesis as described elsewhere (34). *E. coli* BL21 endo^-^ cells transformed with pPRObLLOC484AD69C plasmid were independently grown in Terrific Broth, and proteins were expressed upon induction with isopropyl thiogalactopyranoside (IPTG). Cells were lysed and proteins were purified using Ni-NTA bead based affinity chromatography as described earlier (5). Purified proteins were desalted in buffer and were then run in SDS-PAGE gel to check for purity. It was then labeled by maleimide-cysteine covalent linkage as reported earlier (35). The purified proteins were quantified using standard Bradford method (36) and the labeling efficiency (LE) was calculated based on the report described earlier (37). The LE of the protein was calculated to be 0.72 and its activity was verified by standard hemolysis. Detailed protocol is provided in *SI Text*.

### Supported lipid bilayer (SLB) preparation

SLBs are prepared using Langmuir-Blodgett (LB) technique as described elsewhere (38). Cleaned and RCA-treated glass coverslips were mounted to Langmuir-Blodgett trough (KSV-Nima) maintained at 15°C. ≈ 50 to 60*μ*l of 1mM lipids dissolved in chloroform are spread on the air-water interface. After the solvent evaporates, the monolayers were compressed at a constant rate to attain a target surface pressure (Π) of 32 mN/m at which the layer is transferred on to the substrate. In order to prepare an asymmetric bilayer with leaflet specific labels, the trough was thoroughly cleaned before spreading fresh lipids.

### Bilayer confocal imaging and area analysis

The bilayers kept in MES buffer at pH 5.5 were imaged using Leica SP5 confocal microscope and the images were analysed using Leica LAS-AF and Image-J software. For imaging, a 63X objective, available with the Leica SP5 system was used. Three lasers with 488 nm (Argon), 590 nm (He-Ne) and 633 nm (He-Ne) wavelengths were used for excitation and the emitted light was detected using photo-multiplier tubes in three channels of required wavelength are used appropriately. Detailed description is provided in *SI Text*.

### Fluorescence Correlation Spectroscopy for measuring lipid diffusion

Time Correlated Single Photon Counts (TCSPC) were collected using the Avalanche Photo Diode (APD) detector with 640 to 704 nm filter and were correlated with the Piqo-quant Symphotime software. The correlation curves were analysed and fitted using

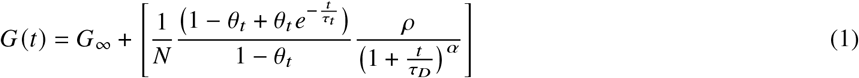

where *G* (*t*) is the correlation function, *G*_∞_ is the offset, *θ_t_* is the fraction that is in the triplet state, *τ_t_* is the time duration to stay in the triplet state, *ρ* is the fraction of diffusing molecules and *τ_D_* is the diffusion time. The Quickfit software was used to obtain the *τ_D_* values from Eq. 1. The diffusion coefficient was evaluated using,

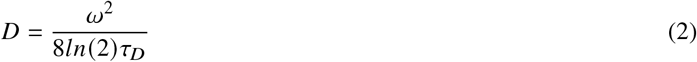

where *ω*=310 nm is the FWHM of confocal beam.

### Förster Resonance Energy Transfer

In order to calculate the efficiency of FRET between Alexa594 (donor) and Atto647N (acceptor), the bilayers treated with known concentrations of LLO, C_*p*_ were imaged initially before bleaching the acceptor with a 633 nm laser. Pre- and post-bleached images were taken using the PMT detectors. Pre- and post-bleached intensities were averaged over 5 frames and the FRET efficiency, *E* was calculated using

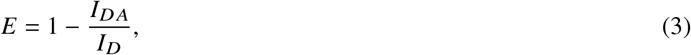

where *I_DA_* is the intensity of donor in the presence of acceptor, *I_D_* is the intensity of donor in the absence of the acceptor. In this case, the FRET occurs from a single donor to multiple acceptor. Hence, the donor and acceptor distance *z* was calculated using,

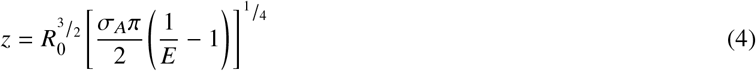

*R*_0_ = 7.089 nm is the Förster distance for Alexa594 and Atto647N pairs calculated as described earlier (39).

### Diffusing Wave Spectroscopy(DWS)

Micro-rheological measurements were carried out using diffusing wave spectroscopy on small unilamellar vesicles (SUVs) with varying LLO concentrations (C_*p*_). The parameters and analysis is described in the *SI Text*.

## RESULTS AND DISCUSSION

### LLO binds to the liquid disordered domain and alters the lipid mobility

The results presented here are based on supported lipid bilayers (SLBs) having lipid composition containing 1,2-dioleoyl-sn-glycero-3-phosphocholine (DOPC), 1,2-dipalmitoyl-sn-glycero-3-phosphocholine (DPPC) and Cholesterol (Chol) in the ratio of 2:2:1, abbreviated as 221, similar to our recent study (8) on GUVs. The SLBs were incubated with LLO, at different concentrations, C_*p*_. The studied SLBs, prepared by Langmuir-Blodgett (LB) technique exhibit well defined phases, commonly referred to as liquid disordered (L_*D*_) and liquid ordered (L_*O*_), as reported in similar systems (8, 40) and is also evident in the confocal fluorescence microscopy images presented in Fig. 1*A*. What is also visible, in the same images is a clear preference of LLO binding towards the L_*D*_ phase (8).

**Figure 1:**
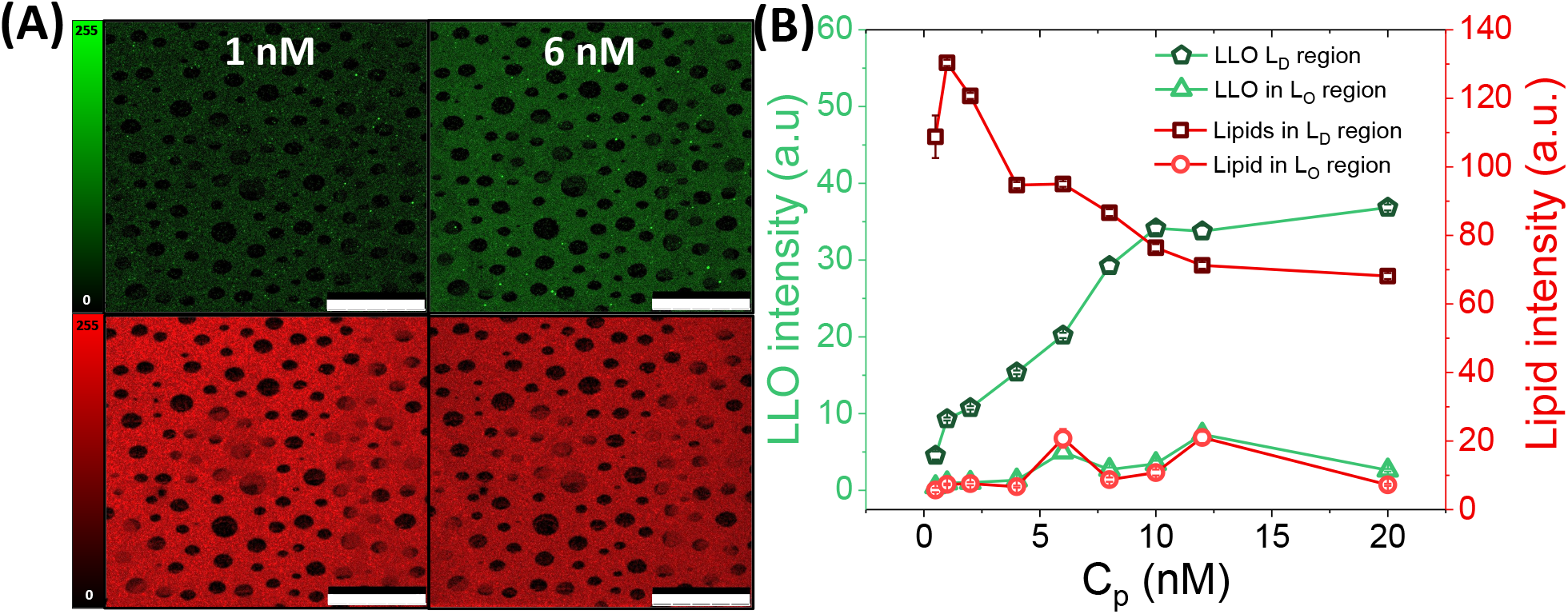
**A.** DOPC:DPPC:Chol (2:2:1) bilayer transferred using the LB method labeled with Atto647NDMPE (red, I_*D*_ specific dye) exposed to 1 nM and 6 nM LLO labeled with Alexa594 (green). **B.** Fluorescence intensity of LLO and lipids on DOPC:DPPC:Chol (2:2:1) plotted as a function of concentration of LLO concentration (C_*p*_) showing initial increase (or) decrease respectively in the L_*D*_ region. The fluorescence intensity in the L_*O*_ region is relatively unchanged.

The fluorescence images reveal a decrease in lipid intensity in the L_*D*_ phase with a concomitant rise in LLO intensity with increasing LLO concentration, C_*p*_ suggesting a strong correlation between protein binding and lipid concentration. Upon quantifying the fluorescence intensity changes, we observed that LLO intensity, initially rises and then saturates at C_*p*_ between 10-12 nM, while the lipid intensity, simultaneously decreases saturating at around similar values of C_*p*_. Notably, the intensity in the L_*O*_ phase remains relatively unchanged over this concentration range suggesting that LLO induced structural remodeling is largely restricted to the L_*D*_ phase. In addition to the phase separated 221 system, we also investigated a membrane system containing cholesterol with 3:1, DOPC:Chol ratio, abbreviated as 31, which exists in the homogeneous one phase regime at room temperature. The trend of increasing LLO intensity (*SI Appendix*, Fig. S4) with increasing C_*p*_, is found to be qualitatively similar to that observed for the 221 system. The decrease in lipid intensity in the L_*D*_ phase in the 221 system is suggestive of phenomenon associated with lipid removal from the bilayer upon exposure to LLO.

While this data provides a clear correlation between increase in LLO concentration and loss of lipids possibly due to pore formation it does not provide quantitative information about the underlying membrane dynamics or mechanical property changes nor does it provide information about the LLO conformational changes in terms of membrane insertion. We now turn our attention to lipid diffusivity, *D*, measured in the L_*D*_ phase as revealed in Fig. 2*A*, by the C_*p*_ dependent confocal fluorescence correlation spectroscopy (FCS) correlation curves. A clear decrease in lipid correlation time, *τ_D_*, with initial increasing C_*p*_ is observable. However, this trend reverses and *τ_D_* starts increasing above a certain value of C_*p*_. To quantify this further, we have extracted *D* values from FCS correlation data collected over multiple regions of the SLBs at each value of C_*p*_ (see methods and Table. S1). This data is presented in Fig. 2*B* where a clear non-monotonic trend is revealed from this data. Interestingly the regime of increasing diffusivities coincides both with the concentrations of C_*p*_ where we observe the maximum rate of decrease in lipid intensity and the concomitant rise in LLO intensity. This suggests a possible link between the membrane reorganization as reflected in the lipid dynamics, LLO oligomerization and pore formation. Fig. 2*C* shows similar FCS correlation data for the 31 system corresponding to different C_*p*_. A similar non-monotonic trend in *τ_D_* is visible. This is further quantified in Fig. 2*D*. The non-monotonic nature is revealed in this system as well although there are quantitative differences in terms of actual values of *D* as well the dynamical cross-over concentration C_*p*_. This points to a common microscopic origin for the observed dynamical cross-over in these lipid membranes induced by LLO while the difference in actual cross-over concentrations C_*p*_ is likely related to the intrinsic fluidity difference between the two lipid compositions as well as to the organisation and availability of cholesterol for LLO pore formation as has been observed earlier (9, 10, 41).

**Figure 2:**
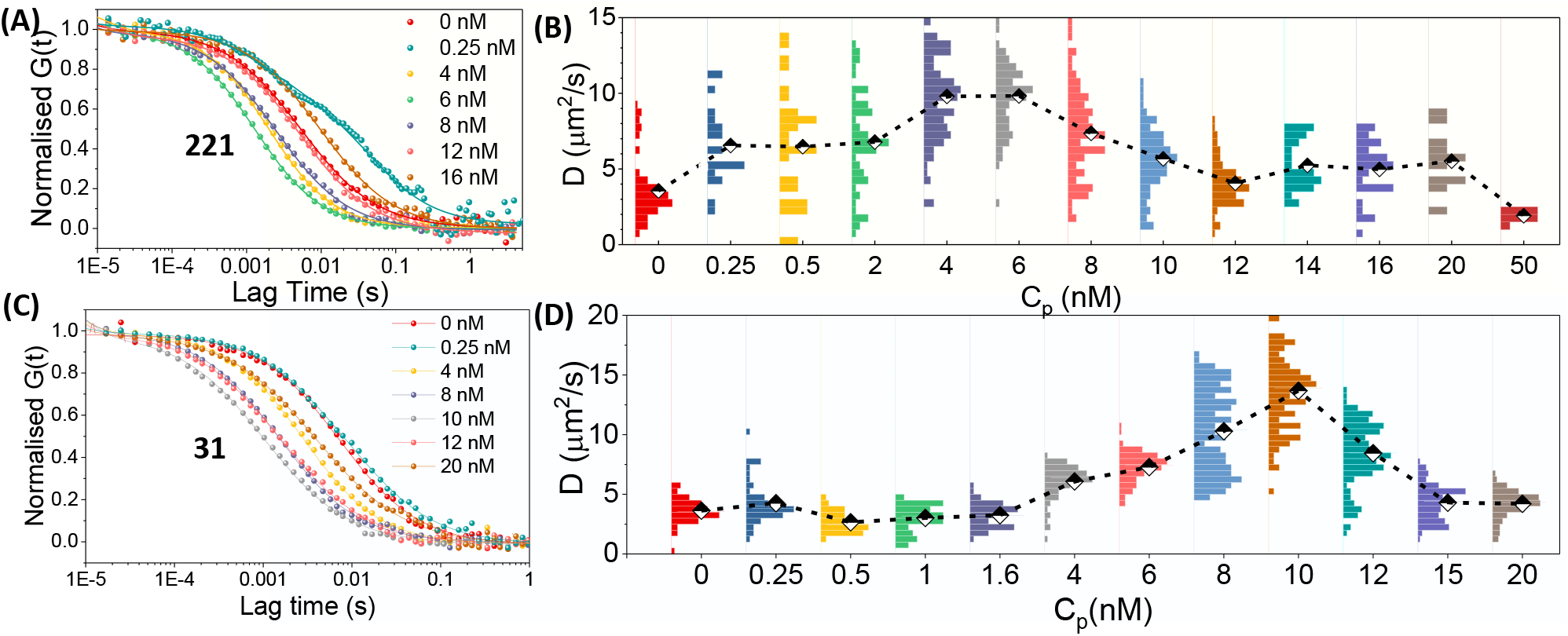
**A.** Normalised FCS autocorrelation curves with corresponding fits for DOPC:DPPC:Chol (221) lipid diffusion in the L_*D*_ domains measured at different LLO concentrations, C_*p*_. **B.** Lipid diffusion coefficients *D* of DOPC:DPPC:Chol (221) calculated from the relaxation time, *τ_D_* (Eq.2) of the correlation curves. The histograms are plotted to illustrate the distribution of *D* across the entire bilayer sample and the dotted lines connect the mean *D* values at each C_*p*_. Corresponding data are shown for the DOPC:Chol (31) system (**C**) and (**D**)

Significantly, all reports of earlier studies of protein, including AMPs, concentration dependence on lipid or protein diffusion on biomembranes only report a decreasing diffusivity, similar to the high C_*p*_ regime observed here (20, 42, 43). In fact with AMPs, which are similar to PFTs, in terms of their pore forming abilities on microbial membranes, lipid diffusivity was found to decrease with increasing concentration (17, 18). However, there are no reports of the increasing D regime with C_*p*_, to the best of our knowledge. While the lipid diffusivity data provides information about membrane response, no quantitative information can be gleaned about membrane bound protein conformational changes and oligomeric state distributions.

### Presence of arcs and pores estimated from free area model

In what follows, we develop a model to explain the observed non-monotonic dependence of lipid *D* on C_*p*_. In order to relate changes in lipid diffusivity, *D* to changes in free area, we consider the extension of the model (44), for lateral lipid diffusion, that occurs in 2-dimensions (2D), developed by Almeida *et.al*.,(30). In this model the relationship between the lipid diffusivity, *D* and the lipid free area, *a_f_* is given by,

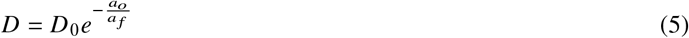

where, *D*_0_ is the reference lipid diffusivity, *a_f_* is the free area per lipid, *a_o_* is the minimum area per lipid above which lipid diffusion occurs. Since our system consists of a mixture of protein and lipids it is more convenient to use an extensive representation for the areas. If *A_t_* represents the total observation area, *A_f_* is the total available free area at a given C_*p*_, and *A_o_* is the minimum area required for diffusion, then

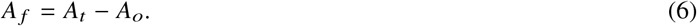

We note that *A_o_* is a function of the protein and lipid concentration at a given C_*p*_. Combining Eq. 5 with Eq. 6 in the extensive area representation we obtain,

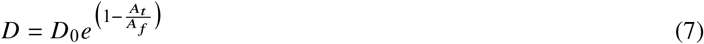

which relates the variations in *D* to *A_t_* and *A_f_*. Using Eqn. 7 we estimated the free area fraction *A_f_* /*A_t_*, by using the lipid *D* values measured from FCS, and taking *D*_0_ = 21 *μ*m2/s (*SI Text*).

At a fixed protein concentration, the total occupied area consists of contributions from both the added LLO (*A_p_*) and lipids (*A_l_*) after membrane remodelling events have occurred due to LLO pore formation. The area occupied by the protein *A_p_* can be obtained from,

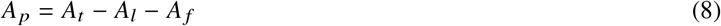

In order to evaluate the area of lipids, *A_l_* we use the mean area per lipid obtained as a weighted average for a given composition from the measured Langmuir isotherms at the air-water interface (Fig. S2). The number of lipids is estimated from the FCS sampling area in the L_*D*_ phase and scaled to the sampling area *A_t_* (detailed analysis provided in *SI Text*). A key observation is the sharp decrease in the area *A_l_* (Fig.S6 *D,E*) as C_*p*_ is increased. This is a distinct quantification of the loss of lipids from the membrane, indicative of pore formation, corroborating the lowering of lipid intensity shown in Fig. 1*B*. Importantly the C_*p*_ at which a minima is observed coincides with the crossover point in the diffusivity data observed in Fig. 2. Furthermore the shift in the crossover point from 221 to the 31 membranes is also captured quantitatively. Using these values of *A_l_* at a fixed value of *A_t_*, with *A_f_* estimated from Eq. 7, *A_p_* is evaluated using Eq. 8 for each value of the protein concentration C_*p*_ as shown in (Fig.S6 *D,E*) where the areas have been scaled by *A_t_*. In contrast, fractional *A_p_* values show a monotonic rise with increasing C_*p*_ consistent with the LLO intensity (Fig. 1) trends in this regime.

Once an estimate of *A_p_* is obtained from Eq 8, the key step in our model is to correlate the variations in the area contributions from LLO to possible oligomeric states on the membrane which evolve as a function of C_*p*_. To simplify and quantify the contribution of these states to changes in free area and hence lipid diffusion, we assume that the predominant oligomeric states consisting of arcs (A) and rings (R) contribute to the LLO intensity at a particular C_*p*_ (Fig. 1). To proceed further we need to quantify the number of oliogmeric units of the different oligomeric states present on the membrane within the area *A_t_* for a specific LLO monomer concentration C_*p*_ in the bulk solution. The number of LLO units present in *A_t_*, at a given C_*p*_ is assumed to be proportional to the total LLO fluorescence intensity of the protein from which the number of LLO oligomers *L_p_* is estimated (detailed analysis provided in Fig S6 *C*) at each value of *C_*p*_*. Although in reality the contributions to *A_p_* arise from a distribution of oligomeric states at a given C_*p*_, we use a simplified 2-state model wherein area contributions arise from a linear combination of arcs and rings (see derivation in *SI Text*). Within this framework,

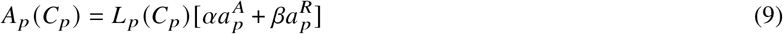

where, 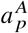 is the mean area per arc, 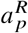 the mean area per ring. Based on the reported LLO hydrodynamic radius of 3.56 nm (10) and mean arc and ring diameters of 31.3 nm and 50 nm (5) respectively, we obtain 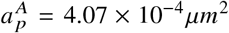, and 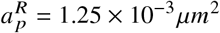. These sizes represent mean values obtained from the distributions observed in AFM experiments (5)

The coefficients *α* and *β* are obtained by solving Eq. 9 (C_*p*_ **>** 2 nm) subject to the normalization condition *α* + *β* =1 (see derivation in *SI Text*). Using this procedure the values of *α* and *β* are obtained exactly (see Table. S2). However for C_*p*_ = 10 and 12 nM (221) we had to impose an additional constraint (0 < *α*, *β* < 1) to obtain meaningful solutions. Using these *α* and *β* values and the corresponding *A_p_* from Eq. 9 we re-evaluate *D* using Eqs. 8 and 7 and compare with *D* values obtained from FCS measurements (Fig 3*A* and *B*) for the 221 and 31 systems respectively. As expected the model predictions are excellent with small deviations observed at C_*p*_ = 10 and 12 nM for the 221 system where additional constraints were imposed.

**Figure 3:**
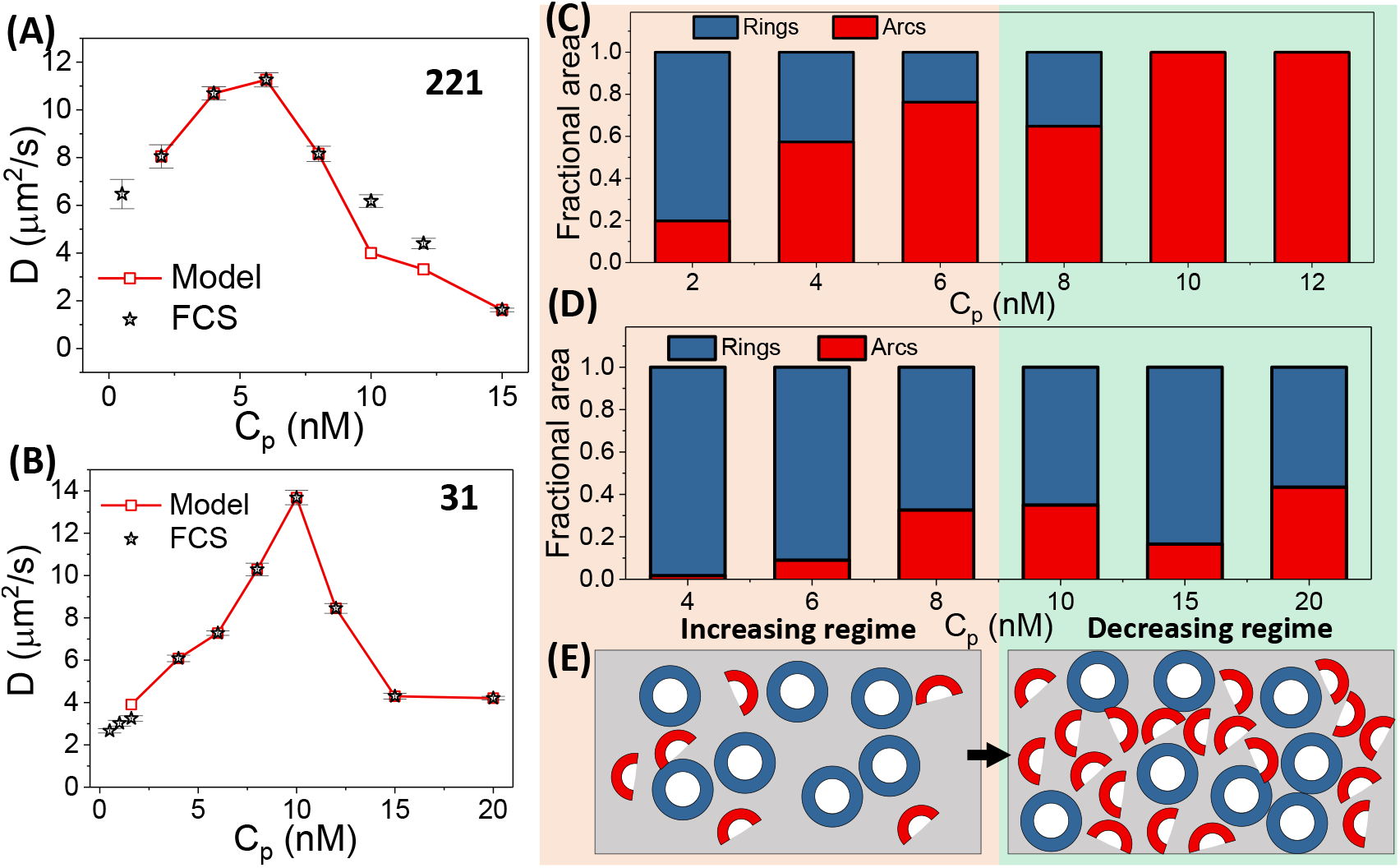
**A.** Lipid diffusion coefficients *D* measured from FCS (black star) on 221 bilayer compared with the *D* values recalculated based on the solution from the proposed two-state model (red). **B.** Data similar to panel **A** provided for the 31 lipid bilayers. Fractional area populations of arcs and rings calculated from the evaluated *α* (arcs) and *β* (ring) values (Table. S2) for the increasing and decreasing diffusivity regimes of DOPC:DPPC:Chol (221) (**C**) and DOPC:Chol (31) (**D**). **E.** Schematic showing the oligomeric distributions for the increasing and decreasing diffusivity regimes.

From the solutions for *α* and *β* we evaluate the fractional area contributions from arcs and rings illustrated in Fig. 3*C*, for the increasing and decreasing diffusivity regimes for the 221 system. Interestingly, we observe a transition in oligomeric states from one dominated by rings at low concentration to arcs at high concentration. This transition correlates with the dynamic transition observed in lipid diffusivities (Fig. 2) as a function of C_*p*_. Presence of rings, which are fully inserted pore states, are clearly favoured to eject lipids (21, 22), is consistent with the expectation of lipid membrane excision driven lipid loss leading to increase in available free area for the membrane lipids. This increase in free area for the lipids should lead to an increase in diffusivity as per the free area models. In contrast to the 221 system the changes in fractional populations of arcs and rings is more subdued for the 31 system (Fig. 3*D*) and we observe that although there is a tendency for an increase in arc population, rings remain the dominant population as a function of C_*p*_. This is probably why we observe slightly higher diffusivity values and a higher dynamical crossover concentration for the 31 system when compared to the 221 system. Significantly, the results suggests a possible connection of the oligomeric state probability to the cholesterol availability for LLO to bind which in turn determines the ease and nature of pore formation (8–10, 45).

However, what is more subtle, is the role played by the preponderant arc like structures beyond the cross-over concentration where lipid diffusivities begin to decrease. It appears that these arcs are not very efficient in ejecting lipids since we do not see much decrease in lipid intensity (Fig. 1*B*) or in number of lipids, (Fig. S6 *A*) suggesting that these oligomeric states lead to a decrease in free area for lipids effectively enhancing crowding effects and a consequent decrease in diffusivity. Indeed molecular dynamics simulations reveal that lipids are displacement from the arc interior into the surrounding lipid milieu (21, 23) While the two state model for membrane bound arcs and rings provides a connection between the relative population of these state and membrane areal changes, the extent of penetration of these oligomers into the membrane is as yet unknown. Recently it has been observed that oligomeric arc and pore states can exist in either a membrane bound or membrane inserted states, discerned from height variations in AFM measurements. (5, 10). In order to obtain information about the extent of membrane insertion of these oligomeric states we resorted to Förster Resonance Energy Transfer (FRET) measurements which are discussed next.

### Structural changes of LLO observed using FRET with varying LLO concentration

From the FRET measurements, the C_*p*_ value at which the distance between the donor-acceptor pair, *z* saturates (Fig. 4 and Fig. S8) corresponds closely with the cross-over concentration for *D*, (Fig. 2) suggesting a correlation between pore forming protein induced membrane structural remodeling, LLO insertion and membrane dynamics. Moreover, it is clear that the oligomeric states that were obtained from the solution of Eq. 9, as discussed earlier, as most probable conformations in the different dynamical regimes are membrane inserted pore complexes and not pre-pores. This explains why the ring-like structures are efficient in lipid ejection leading to increase in free area for remaining lipids in their respective membrane systems.

**Figure 4:**
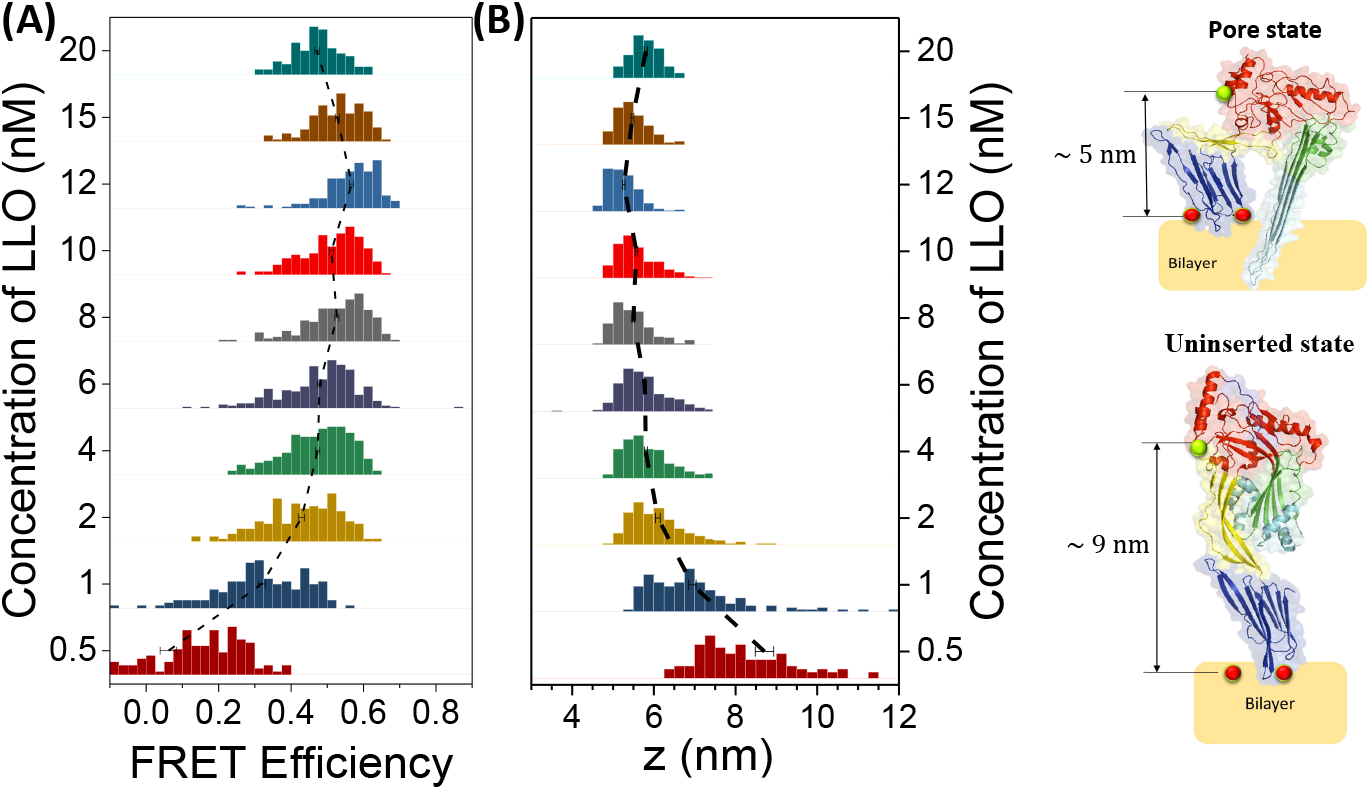
**A.** FRET efficiency measured using Eq. 3 between Alexa594LLO (donor) and Atto647N (acceptor) of DOPC:DPPC:Chol (221). During FRET measurements, the lipids were labeled only on the upper leaflet and data is collected only from L_*D*_ domain of the bilayer. **B.** Distance between the donor-acceptor pair calculated based on Eq. 4 denoted as ‘*z*’. The data reveals populations of both inserted pore states and uninserted pre-pore states in the lower concentrations shifting to a population of completely inserted pore states above 2 nM. Images inside panel **B** depict the structure of single LLO monomer in its pre-pore state and inserted pore state. Note that the structures are representative images and LLO monomers do not insert prior to oligomerisation.

### Rheological properties of vesicles observed with varying LLO concentrations

Since our diffusivity data as well as inferences in the changes in free area for lipids suggest active membrane remodelling during pore formation by LLO, we expect these dynamical changes to reflect in macroscopic rheological or mechanical properties of the membrane. Furthermore, the changes in mechanical properties can also be probed at the cellular level to reflect large scale membrane modulations upon exposure to PFTs. To address this question we performed diffusing wave spectroscopy (DWS) based micro-rheology (MR) measurements on small unilamellar vesicles (SUVs) of similar composition DOPC:DPPC:Chol (221) as used in the SLBs at 24°C.

The auto-correlated and normalized scattered intensity changes with time is illustrated in Fig. 5*A*. Careful scrutiny of the data also suggests a non-monotonic behaviour of the correlation time with C_*p*_. The data was further analysed (46) to extract the MR parameter, *G”*, for all the data as shown in Fig. 5*B*. Interestingly, we again see a non-monotonic trend of *G”* with C_*p*_, which mirrors the variation of *D*. This also lends credibility to the pore formation driven lipid ejection induced increase in lipid free area which in turn has a tendency to fluidize the membrane leading to a significant reduction of *G”* for the SUVs. However, the observed increase of *G”* at higher concentration, post the saturation of the pore formation process suggests a very interesting self-stabilizing mechanism of the SUVs leading to an overall stiffening of the membrane which takes place in a crowded regime dominate by membrane arcs (Fig 3).

**Figure 5:**
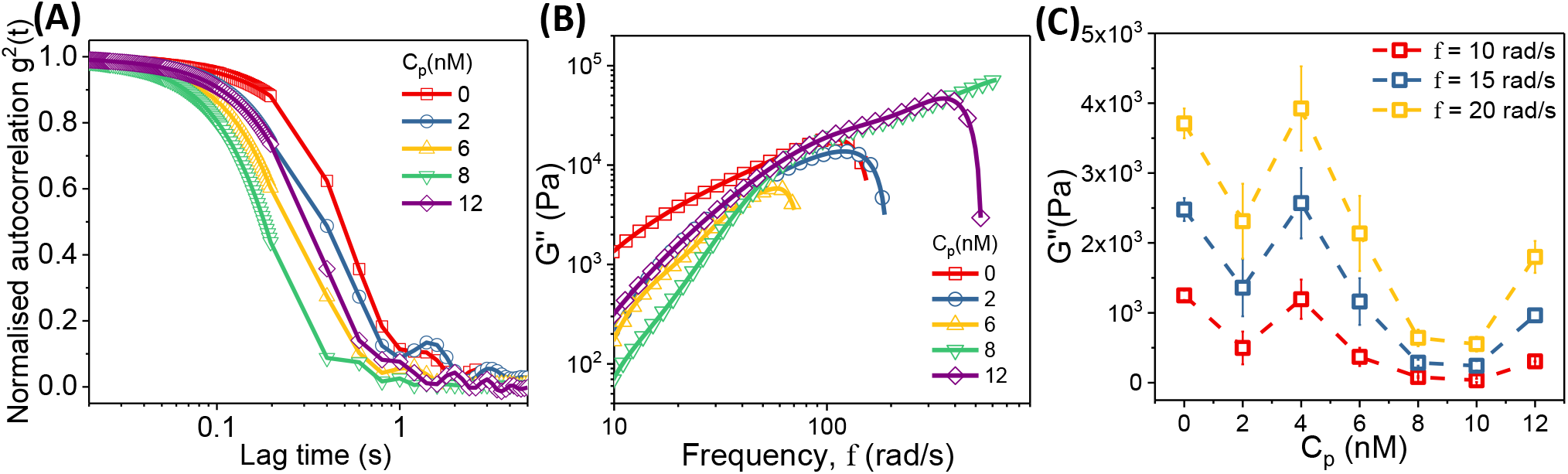
Diffusing wave spectroscopic data for the small unilamellar vesicles as a function of LLO concentration, C_*p*_, measured at 24°C. The normalised autocorrelation *g*^2^(*t*) (**A**) and corresponding loss moduli, *G”* (**B**) values provided indicate differences in the rheological properties with respect to C_*p*_. **C.** The loss moduli, *G”* provided as a function of C_*p*_ at a given frequency, f.

## CONCLUSION

Despite the knowledge about the pore formation pathways, membrane remodelling and related repair processes during attack by PFTs remain largely unexplored. Here we systematically investigate membrane remodelling events as reflected in lipid dynamics in membranes exposed to LLO over a broad range of protein concentrations to provide a hitherto unexplored link between lipid ejection and membrane inserted pore states. Significantly enhanced lipid mobility is linked to the formation of ring-like pores at intermediate protein concentrations where lipid ejection plays a dominant role. What is most interesting is the existence of a cross-over concentration at which the ring-like pore formation process, possibly, reaches saturation and hence lipid ejection becomes less significant while lipids increasingly feel the effect of crowding due to arc-like pore complex formation which, although fully inserted, are not efficient in lipid ejection. While it is possible that both the mechanisms (free area creation due to lipid ejection and pore induced crowding) co-exists over a certain concentration range the clear separation between the two regimes that emerges suggests dominance of one type of oligomeric state over the other in these two regimes.

Our proposed simplified two state model with contributions from arcs and rings is sufficient to capture changes in lipid free area variations predicted using the free area model developed by Almeida *et al*., (30). The model tacitly assumes that oligomeric states which contribute to changes in lipid available free area are membrane inserted, thereby excluding populations of membrane bound monomers or pre-pore states whose contributions to lateral changes in lipid area are expected to be minimal. This presence of membrane inserted oligomeric populations is supported by FRET data which reveals the increase in membrane inserted states with increasing C_*p*_. Interestingly, the slight decrease in FRET efficiency, and a consequent increase in *z*, at high C_*p*_ values (> 12 nM) in the 221 membranes correlates with the crowded regime of lowered *D* values. This suggests emergence of a sub-population of oligomeric states with decreased penetration in the membrane when compared with fully inserted states with lowered *z* values (Fig. 4). In this regime, the 2-state model predicts a preponderance of arcs (Fig. 3) and recent molecular dynamics simulations reveal that arc-like states have a tendency to protrude outward from the membrane plane when compared with the fully formed ring-like pore states (21, 47) We note that the regime of low values of C_*p*_ ( < 1 nM) lies outside the purview of the two-state model, where LLO is more likely to sample membrane bound non-inserted states.

The active membrane remodelling processes captured in our study has implications in several calcium signalling driven cellular repair pathways in order to mitigate cell lysis and recover membrane and cellular integrity. The repair processes involve active or passive membrane remodelling events such as exocytosis and endocytosis in order to rid the membrane of the damaged toxin associated sites on the plasma membrane (11, 48–50). Our study suggests that these membrane repair processes are more likely to occur at intermediate protein concentrations where lipid mobility is higher. Our results could lead to developing effective strategies for preventing cell lysis induced by virulent pathogen PFT attack and hence mitigation of severe infections and deadly diseases.

## AUTHOR CONTRIBUTIONS

The authors thank Department of Science and Technology - Science and Engineering Research Board (DST-SERB) for funding the project. IIP thanks Rudradeep for discussions regarding the model, Pradeep Satyanarayana for assistance in preparing labeled LLO and Sandhya S. Visweswariah for providing the plasmid as well as assistance with protein purification.

## ACKNOWLEDGMENTS

We thank G. Harrison, B. Harper, and J. Doe for their help.

## SUPPLEMENTARY MATERIAL

An online supplement to this article can be found by visiting BJ Online at http://www.biophysj.org.

